# Novel, economically important semi-dwarf and early mutants: Selection and development from Improved White Ponni Rice (*Oryza sativa* L.)

**DOI:** 10.1101/500637

**Authors:** S. Ramchander, Andrew Peter Leon, Jesima Khan Yasin, KK. Vinod, M. Arumugam Pillai

**Affiliations:** Visiting Scientist (SERB –National Post-Doctoral Fellow), IRRI-South Asia Hub, ICRISAT, Patancheru, Hyderabad, India; Department of Plant Breeding and Genetics, Agricultural College and Research Institute, Killikulam Tamil Nadu Agricultural University, India; Scientist, Division of Genomic Resources, ICAR-National Bureau of Plant Genetic Resources, and Faculty, ICAR-IASRI, New Delhi, India; Principal Scientist Rice Breeding and genetics research Centre, Aduthurai 612 101 Tamil Nadu, India

**Keywords:** Early maturing, irradiation, LD_50_, lodging resistance, semi-dwarf, Single Marker

## Abstract

Rice variety, Improved White Ponni is a medium duration crop, but highly susceptible to lodging impacting maximum yield losses. The present investigation aimed to identify early and early semi-dwarf mutants in Improved White Ponni by inducing variations using gamma rays without altering its native grain quality traits. Seeds were treated with various doses of after fixation of the LD_50_ value of gamma radiation and reported that most of the traits exhibit variations in the mutants at various levels of irradiation. The selection for earliness and dwarf plant height was imposed in M_2_ and it was confirmed by evaluation of M_3_ generation. Apart from semi-dwarf early mutants, high tillering habit, narrow rolled leaf, upper albino leaf and grassy stunt and extreme dwarf mutants were also identified. Characterization of mutants using already reported genic and linked microsatellite markers associated with semi-dwarfism and earliness resulted that PIC value ranges between 0.037 and 0.949 with an average of 0.382. Single marker analysis revealed that RM302 and RM310 on chromosome 1 and 8 had exhibited an association with the traits plant height, culm and internodal lengths. Of these gene-specific markers, GA20Oxi_1 and GA20Oxi_2 have shown polymorphism among mutants. GA20Oxi_2 showed null alleles in the dwarf mutants and this clearly emphasized that there are some base deletions exists in the region of exon 2 of *sd1* region. GA_3_ response study shown that identified mutants were GA_3_ responsive except IWP 11-2, IWP 48-2, IWP 50-11 and IWP 33-2 which showed very low responsive. Agar plate assay revealed that, IWP 50-11, IWP 33-2, IWP 43-1, IWP 47-2 and IWP 18-1 had low production of α- amylase. Scanning electron microscope examination on confirmed mutants exhibited larger cell size and a lesser number of cells per unit area than the wild-type which shows that these mutants are defective in GA mediated pathway.

## Introduction

Rice (*Oryza sativa* L.) improvements especially in case of quantity and quality enhance the rice global production and play a vital role to overcome food shortage, enlighten the local consumption and export. Introducing new varieties of rice characterized by early heading, short stature, lodging resistance, blast resistance and improved grain quality characters are the main objectives for a quantum increase of grain yield of rice. Semi-dwarfism, an important trait in cereals and governs by green revolution gene *semi-dwarf 1* (*sd-1*), which have an. impact of short and thick culms, imparts lodging resistance and nitrogen responsiveness and it has initially derived from Dee-Gee-Woo-Gen and responsible for higher yield without affecting the native grain quality parameters of the variety (Futsuhara and Kikuchi, 1997). Kikuchi et al. (1985), identified several *sd-1* mutants in rice and these mutants has been used in several breeding programs. It was found that *sd-1* gene was responsible for the production GA20 oxidase-2 enzyme involved in the catalytic steps of gibberellin (GA) bio-synthesis (Spielmeyer *et al.*, 2002; Sasaki *et al.*, 2002;). A defect in the production of GA was one of the key determinants for semidwarf plant type in most of the *sd-1 mutants* (Sakamoto *et al.*, 2004) caused by low GA production due to varied (either loss or reduced) function of GA20 oxidase-2. In *indica* variety IR8, an allelic form of *sd-1* contains 382 bp deletions from exonic regions of *sd-1* locus of 1 to 2 resulting in formation of stop codon, which ultimately modifies the gene function. Whereas in the cases of some *japonica* varieties namely Jikkoku, Calrose76and Reimei, single base substitutions lead to a single amino acid change in the *sd-1* gene (Spielmeyer *et al.*, 2002; Ashikari *et al.*, 2002).

Rice, as a facultative short-day plant and early heading or flowering, has a significant impact on the regional adaptability of the rice varieties. Number of QTLs has been found in crosses among wild strains of rice, *japonica* and *indica* strains. Apart from genetic analysis, induced mutations played a pivotal role for improving rice architecture by developing a large number of variants such as, early, dwarf, high tillering, blast resistance, low amylose and high yielding mutants (Soomro *et al*, 2006). The basic requirement for direct improvement target agronomic trait, available genetic variability is required to meet the demand of the breeder. Therefore, induced mutations with the discovery of an array of radiation mutagen and improved treatments methods offer the possibility for the induction of desired changes in various attributes, which can be exploited as such or through recombination breeding (Cheema and Atta 2003). Hence, the primary objective of this study is to induce variations in Improved White Ponni (IWP) by using gamma irradiation and to develop desirable semi-dwarf, early high yielding mutants with improved grain quality parameters.

## Materials and Methods

### Genetic material

Improved White Ponni, an important medium duration (115 days to flowering) and quality rice variety in southern parts of India for its fine slender grains but had a problem of tallness (> 150 cm) which make the crop susceptible to lodging and grain loss. The seeds of Improved White Ponni were subjected to gamma irradiation at different doses (100Gy to 500Gy) by using Gamma Chamber – Model GC 1200 installed at Tamil Nadu Agricultural University, Coimbatore. The experimental plots were raised at Agricultural College and Research Institute (Killikulam) and Agricultural Research Station (Thirupathisaram) during the year 2011 to 2014.

### Mutagenic treatment, selection and evaluation

Five hundred well-filled seeds of IWP were treated with gamma rays at various doses from 100 to 500 Gy with an interval of 100Gy. After treatment, M1 seeds were immediately sown in raised nursery beds along with control seeds. On 25^th^ day after germination, the seedlings were planted in the main field where standard cultural practices were followed and harvested on single plant basis (M2 seeds). The M_2_ generation was raised from individual M_1_ plant following plant to progeny method. A total set around 184 M_1_ (families) plants seeds were forwarded to M_2_ generation whih was raised during *rabi* 2012 to summer 2012 (September 2012 to April 2013) without replication. The selection was imposed for dwarf plant type and earliness in flowering along with other desirable characters. A set of 152 mutants were identified in M_2_ and forwarded to M_3_ generation for their evaluation and validation. These mutants were sown in raised nursery beds and transplanted to the main field 28 days after sowing in three replications during *Kharif* 2013 (May 2013 to November 2013). M_3_ generation was evaluated for various traits associated with the trait of interest, yield component traits and traits controlling quality parameters fetching higher consumer’s preference by the methods given in International Rice Research Institute, Standard Evaluation System (SES) 2013. Estimation of amylose content of identified mutants was carried out by calorimetric method ((Juliano, 1979).

### Study of GA_3_ response in identified mutants

Mutant seeds of IWP along with control were surface sterilized for 30 minutes using with 2 per cent HgCl_2_ (Mercuric chloride) and washed with sterile water. After that, seeds were placed over wet filter paper for two days under dark at 30°C for proper germination. Ten uniformly germinated seeds were placed on the plate containing 1 per cent concentration of agar and kept at 25°C for hastening the proper emergence of the second leaf sheath. Then, 1 μl of GA_3_ solution containing 10 mg/ml was dropped to the coleoptile region by using micropipette on the rice seedlings. The length of the second-leaf sheath of five seedlings was measured after five days of treatment to calculate GA_3_ response (GAR) (Murakami, 1968).

### α- amylase activity assay

The embryo-less half seed of the IWP Mutant and control seeds were surface sterilized using 2 per cent HgCl_2_ for 15 minutes, washed using sterile distilled water for five minutes and placed perpendicularly on plates containing 2 per cent agar medium containing soluble starch (0.2 per cent), sodium acetate (10 mM) and CaCl_2_ (2 mM) with the pH of 5.3. One micromole of GA_3_ was added after autoclaving and incubated for 3 days in the dark at 30°C. After incubation, plates were flooded with I_2_-KI (0.72 g/l I_2_ + 7.2 g/l KI) solution in 0.2 N HCl. Transparent halos around the seed was noticed and it gives an indication of production of α-amylase, which results in the digestion of starch in the plat (Lanahan and Ho, 1988).

### Scanning electron microscope (SEM)

Transverse sections of leaf, nodal region and intermodal regions of the unique mutants (Dwarf mutant, narrow rolled leaf mutants and control) were studied for their difference in internal cell arrangement patterns under Scanning Electron Microscope (SEM) (Model: FEI quanta 200 SEM) built in the Department of Nano Science and Technology, Tamil Nadu Agricultural University, Coimbatore.

### PCR amplification and Electrophoresis

A set of 55 microsatellite markers associated with semi-dwarfism and earliness were used to characterize mutants in the M_3_ generation (Table 1). The PCR amplification was carried out using thermal cycler (Applied Biosystems/BioRad) and amplified products were separated by agarose gel matrix (1.5%) containing 1X Tris-Borate EDTA and electrophoresed at 80 volts for 2 hours and visualized with the help of gel documentation system (BioRad).

**Table 1.** List of microsatellite markers used in this study

| S.No. | Name of the primer | Chrom. | Position (cM) | QTL associated | References |
| --- | --- | --- | --- | --- | --- |
| 1. | RM580 | 1 | 68.2 | *qPH-1, qLFI-1* | Feng-hua *et al.*, 2005 |
| 2. | RM246 | 1 | 115.2 | *qPH1.1, qGP-1* | Lincoln *et al.*, 2017, Rabiei *et al.*, 2007 |
| 3. | RM543 | 1 | 145.6 | *qHd1* | Rabiei, 2007 |
| 4. | RM302 | 1 | 147.8 | *qTGW-1, qDFF1-1* | Sangodele *et al.*, 2014, Marathi *et al.*, 2012 |
| 5. | GA20 Oxi_1 | 1 | 147.8 | *Sd-1* | Spielmeier *et al.*, 2002 |
| 6. | GA20 Oxi_2 | 1 | 147.8 | *Sd-1* | Spielmeier *et al.*, 2002 |
| 7. | GA20 Oxi_3 | 1 | 147.8 | *Sd-1* | Spielmeier *et al.*, 2002 |
| 8. | RM263 | 2 | 127.5 | *qPh2.2* | Lin *et al.*, 2011 |
| 9. | RM250 | 2 | 170.1 | *qPh2.2* | Lin *et al.*, 2011 |
| 10. | RM5430 | 2 | 111.5 | *wt100-vb2.1* | Susanto *et al.*, 2008 |
| 11. | RM5699 | 2 | 42.1 | *qHPH2d* | Zhao *et al.*, 2016, |
| 12. | RM240 | 2 | 158 | *qHd2* | Lin *et al.*, 2011 |
| 13. | RM207 | 2 | 191.2 | *qHd2* | Lin *et al.*, 2011 |
| 14. | RM5849 | 3 | 18.4 | *qHA3-3* | Takeuchi *et al.*, 2008 |
| 15. | RM7365 | 3 | 49.3 | *qTGW3-1* | Hori *et al.*, 2012 |
| 16. | RM545 | 3 | 35.3 | *qHD-3* | Liu *et al.*, 2012, Lin *et al.*, 2011 |
| 17. | RM7 | 3 | 64 | *qDTH3.1* | Moncada *et al.*, 2001 |
| 18. | RM3203 | 3 | 2.2 | *qHd3* | Lin *et al.*, 2011 |
| 19. | RM252 | 4 | 99 | *qPh4* | Lin *et al.*, 2011 |
| 20. | RM567 | 4 | 153.6 | *qPh4* | Lin *et al.*, 2011 |
| 21. | RM5709 | 4 | 109.9 | *qPHT4-3* | Marathi *et al.*, 2012 |
| 22. | RM551 | 4 | 0 | *qPHT4-1* | Marathi *et al.*, 2012 |
| 23. | RM430 | 5 | 78.7 | *qPh5* | Lin *et al.*, 2011 |
| 24. | RM480 | 5 | 130.6 | *qPh5* | Lin *et al.*, 2011 |
| 25. | RM541 | 6 | 75.5 | *qPh6.1* | Lin *et al.*, 2011 |
| 26. | RM30 | 6 | 125.4 | *qPh6.1* | Lin *et al.*, 2011 |
| 27. | RM253 | 6 | 37.0 | *qHD6.1* | Zhao *et al.*, 2016 |
| 28. | RM7023 | 6 | 51.3 | *qGT6d* | Lapitan *et al.*, 2009 |
| S.No. | Name of the primer | Chrom. | Position (cM) | QTL associated | References |

### Statistical analysis

The estimation of mean, analysis of variance and standard error of the traits were worked out by adopting the standard methods (Panse and Sukhatme, 1961). Analysis of phenotypic, genotypic variances and heritability was formulated by Lush (1940), variability parameters like PCV and GCV was determined by theformula given by Burton (1952). Estimation of genetic advance and correlation coefficient among the studied traits was given by Johnson *et al.* (1955). Polymorphism information content (PIC) value of each SSR marker was estimated by using marker scoring data (Anderson *et al.* 1993). The Marker-Trait association analysis between marker data and traits was carried out using Simple linear regression analysis (SLRA).

## Results

### LD_50_ Determination

Probit analysis was carried out using seed germination values to determine the Lethal Dose (LD_50_) of gamma radiation against Improved White Ponni. The expected LD_50_ value for gamma rays in Improved White Ponni was 354.8 0 Gy. M1 plants were harvested individually in all the treatments and used for next season crop (M_2_) where variability was studied for most of the morphological traits.

### Variability for quantitative traits in M_2_ and M_3_ population

#### M_2_ population

In the M_2_ generation, the mean days to fifty per cent flowering ranged from 106.23 (100 Gy) to 112.87 (400 Gy) days whereas wildtype recorded 111.50 days (Table 2a). The lowest mean value of plant height was recorded by 300 Gy (138.68 cm) and the highest mean value of plant height was recorded by 100 Gy (153.43 cm) and these two values were lesser when compared to the mean value of wildtype (162.39 cm). The maximum panicle length was recorded in 400 Gy (25.08 cm) which was followed by 300 Gy (24.86 cm) and 200 Gy (23.73 cm) and the lowest value was noticed in 100 Gy (23.27 cm). A number of grains per panicle ranged from 179.47 (400 Gy) to 197.43 (100 Gy). Primary culm length was ranged from 113.82 cm (300 Gy) to 130.16 cm (100 Gy) and this was less when compared to wildtype (138.14 cm). The trait secondary culm length ranged from 108.52 cm (300 Gy) to 121.37 (100 Gy). The trait thousand grain weight ranged from 16.52 gram (300 Gy) to 16.64 gram (100 Gy and 400 Gy). Single plant yield ranged from 41.38 gram (300 Gy) to 51.11 gram (200 Gy).

**Table 2a.** Mean, variability and heritability estimates of morphological traits in the M2 generation of improved white ponni generated by gamma irradiation

#### M_3_ population

In gamma irradiated population of Improved White Ponni, the mean days to fifty per cent flowering ranged from 104.12 (100 Gy) to 114.26 (200 Gy). The treatment 200 Gy had recorded the highest PCV (9.92), GCV (9.84) and GA as per cent of the mean value of 20.09 with the heritability percentage of 98.32 and this was followed by 100 Gy with heritability and the genetic advance of 99.26 and 17.80 per cent (Table 2b). The lowest plant height was recorded by 100 Gy (113.65 cm) and the highest mean value of plant height was recorded by 300 Gy (126.94 cm) and these two values were lesser when compared to the mean value of wildtype (142.59 cm). The treatment 100 Gy had recorded the highest PCV (15.60), GCV (15.53), heritability (99.14) and genetic advance (31.86). The highest mean value of panicle length was recorded in 400 Gy (26.17 cm) which was followed by 300 Gy (25.73 cm) and 100 Gy (25.02) and these values were higher than IWP (24.92 cm). PCV (12.25), GCV (11.15), heritability (82.85) and GA% of the mean (20.21) was found to be higher in 200 Gy when compared to all other treatments. The trait number of productive tillers per plant ranged from 10.13 (400 Gy) to 16.04 (100 Gy). The treatment 300 Gy had registered the highest PCV (31.05), GCV (28.36) and GA% of the mean (53.36) whilst heritability (86.52) was higher in 100 Gy.

**Table 2b.** Mean, variability and heritability estimates of morphological traits in the M3 generation of improved white ponni generated by gamma irradiation

A number of grains per panicle ranged from 169.70 (400 Gy) to 191.15 (300 Gy). The treatment 100 Gy registered higher value of PCV (14.80), GCV (13.64), heritability (84.92) and GA% of the mean (25.90). The trait primary culm length ranged from 88.77 CM (100 Gy) to 101.52 cm (300 Gy) and this was less when compared to wild-type (116.54 cm). The phenotypic coefficient of variance (18.59), the genotypic coefficient of variation (18.47), heritability (98.70) and GA% of the mean (37.80) was registered higher in 100 Gy. Secondary culm length ranged from 83.93 cm (100 Gy) to 97.80 (300 Gy). First internodal length ranged from 30.33 cm (200 Gy) to 41.52 (100 Gy) and this was lesser than wildtype (43.56 cm). PCV (23.02), GCV (22.47), heritability (95.28) and GA% of the mean (45.19) were noticed to be higher in 200 Gy. The 2^nd^ internodal length ranged from 20.21 cm (200 Gy) to 26.27 cm (100 Gy). The treatment 100 Gy registered the highest PCV (26.76), GCV (25.83), heritability (93.19) and GA% of the mean (51.36). 3^rd^ internodal length ranged from 11.57 cm (100 Gy) to 16.57 (400 Gy). PCV (32.91), GCV (31.16) and GA% of the mean (60.76) were registered higher in 200 Gy whereas heritability (91.48) was found to be higher in 100 Gy. The 4^th^ internodal length ranged from 5.77 cm (200 Gy) to 8.11 cm (300 Gy). The treatment 200 Gy recorded the highest PCV (35.15), GCV (32.87) and GA% of the mean (63.34) whereas the heritability (88.68) was higher in 100 Gy. The trait thousand grain weight ranged from 16.32 gram (100 Gy) to 16.70 gram (400 Gy). The treatment 300 Gy had recorded the highest PCV (1.53), GCV (1.35), heritability (77.63) and GA% of the mean (2.45). Single plant yield ranged from 33.14 gram (200 Gy) to 44.04 gram (400 Gy. The treatment 200 Gy had recorded the highest PCV value of 30.22 whereas other parameters were higher in 100 Gy.

### Identification of unique mutants for agronomic traits

All the mutants were raised under field conditions and screened for novel altered phenotypes in the morphological traits *viz*., flowering, plant height, tillering habit, narrow rolled leaf, upper albino leaf, grassy and extreme dwarf, lodging resistant, lanky culm, *etc*., (Table 3) (Fig 1). Early flowering mutants had registered 79 to 92 days for days to flowering when compared to wild-type which had recorded 112 days to flower. Semi-dwarf mutants identified in this study were early in flowering. High tillering mutants were identified which possess 32 to 44 productive tillers per plant whereas Improved White Ponni had only 18 productive tillers. Narrowly rolled leaf mutants had less panicle length and number of grains with fine grains when compared to wildtype genotype.

**Table 3.** Mutants exhibiting altered morphological traits observed in M2 and M3 generation (gamma rays) of improved white ponni

**Figure1.**
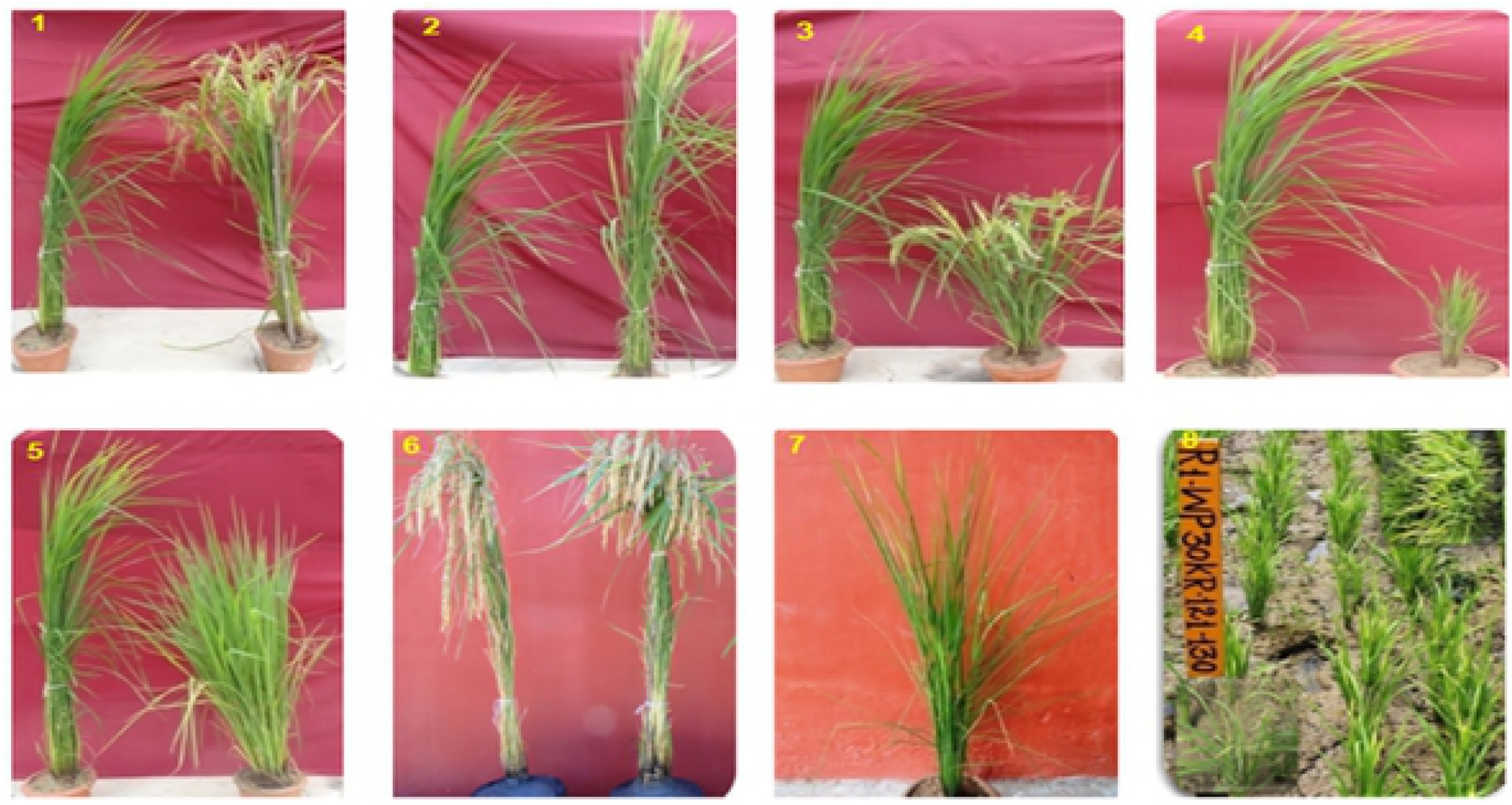
Viable morphological mutants observed in the M_2_ generation of Improved White Ponni 1) Early mutant 2) Tall early mutant 3) Early Semi-dwarf mutant 4) Extremely dwarf mutant 5) High tillering late mutant 6) High tillering mutant 7) Narrow rolled leaf mutant 8) Upper albino mutant

### Morphological and quality trait evaluation of desirable semi-dwarf and early mutants in the M_3_ generation

The ANOVA was computed for all the component traits studied (Table 4) and in case of days to fifty per cent flowering, IWP E-4-3 had recorded lower days of 86.00 and IWP 11-2 recorder higher days of 119.00 days to flower with mean value of 96.93 days. With respect to plant height, the range of 82.21 cm (IWP 59-1) to 142.39 cm (IWP 50-4) was noticed with the mean of 111.18, whereas wildtype had the plant height of 143.26 cm (Fig 2). The trait panicle length exhibited the range of 20.46 (IWP 48-2) to 28.69 (IWP 18-1) with the mean value of 24.17 cm. The trait number of productive tillers ranged from 8.50 (IWP D-1) and 25.50 (IWP 1-12) with a mean value of 13.71. Among the 30 mutants studied, the highest primary culm length was found to be low in IWP 59-1 (59.50 cm) and high in IWP 50-4 (117.33 cm) whereas wildtype had registered higher primary culm length of 114.86 cm. The trait secondary culm length had recorder the area of 83.79 cm with the range of 53.54 (IWP 48-4) to 109.74 (IWP 50-4). With respect to 1^st^ internodal length, the mutant IWP 1-9 had recorded a lower value of 16.63 cm and IWP 51-4 had registered higher value 42.08 cm. The trait 2^nd^ internodal length ranged from 11.54 (IWP 48-4) to 40.06 (IWP 1-9) with the mean of 21.17 cm. In case of the trait 3^rd^ internodal length, the minimum and maximum length were found to be reported in IWP 48-3 (4.77 cm) and IWP 1-9 (21.72 cm) whereas wildtype genotype reported 11.50 cm. The trait 4^th^ internodal length had registered 5.78 cm as a mean with the range of 3.33 (IWP E-4-1) to 10.69 (IWP 1-9) (Fig 3). A number of grains per panicle ranged from 156.00 (IWP 31-2) to 272.00 (IWP 7-1) with the grand mean of 206.07. The trait thousand grain weight was found to be high in IWP 1-9 (21.34 gram) and low in IWP 7-1 (13.26 gram) with the mean of 17.04 gram. In case of single plant yield, the mutants IWP 43-1and IWP 50-4 had recorded higher and lower yield of 46.46 and 25.75 gram, respectively.

**Table 4.** Mean performance of semi-dwarf and early mutants of improved white ponni in M3 generation

**Figure 2:**
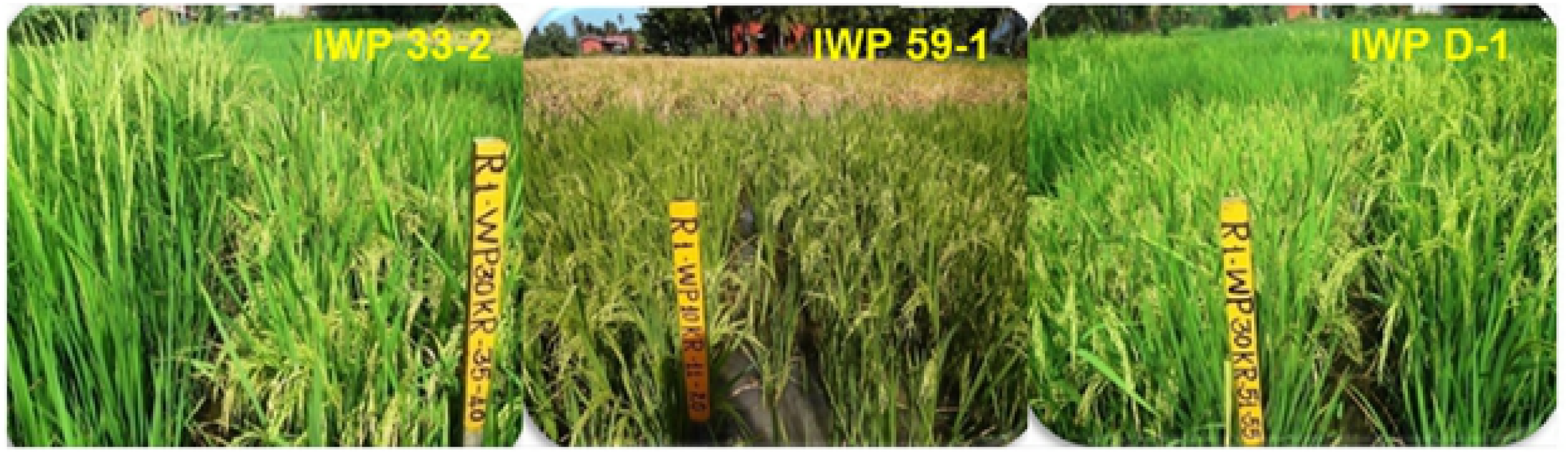
Variation on Morphological traits in identified mutants in M3 generation Flowering and plant height variation Variation on culm length - 1^st^, 2^nd^, 3^rd^ and 4^th^ Intermodal length 1) Improved White Ponni 2) IWP 43-1 3) IWP 59-1 and 4) IWP 50-11

Apart from morphological parameters, this study also involved accessing the quality performance of desirable mutants. The trait kernel length before cooking exhibited maximum in IWP E-2 (6.30 mm) and minimum in IWP 11-1 (4.87 mm) whereas wildtype registered the trait value of 5.58 mm. Kernel breadth before cooking had ranged froM_1_.83 mm (IWP 48-2 and IWP 7-1) to 2.20 mm (IWP 1-9) with the mean of 1.9 mm. With respect to the L/B ratio, the minimum and maximum ratio were found to be observed in IWP 47-2 (2.49), IWP 33-2 (2.49) and IWP 16-6 (3.09). The trait kernel length after cooking revealed that, the mean value of 6.90 with the range of 5.60 mm (IWP D-1) to 7.70 mm (IWP 1-1). In case of kernel breadth after cooking, the minimum and maximum value was found to be noticed in IWP D-1 (2.55 mm) and IWP 1-9 (3.35 mm) with the mean value of 2.93 mm. The trait L/B ratio after cooking showed the mean value of 2.36 with the range of 1.94 (IWP 33-2) to 2.75 (IWP E-3) whereas wildtype genotype recorded 2.37. The maximum and minimum linear elongation ratio was observed in IWP E-3 (1.51) and IWP D-1 (0.99) with the wildtype value of 1.20. The trait BER ranges from 1.28 (IWP 51-4) to 1.69 (IWP 48-2). The maximum and minimum amylose content was found to be noticed in IWP 48-3 (21.26%) and IWP 43-1 (10.40 %) with the mean value of 16.26 %. Gel consistency was ranged from 56.25 mm to 81.25 mm in Improved White Ponni mutants.

### Association analysis

Association analysis found out the relationship among the traits studied and reported that plant height had significantly higher positive association with the traits *viz*., panicle length (0.480), primary culm length (0.970), secondary culm length (0.959), 1^st^ internodal length (0.713), 2^nd^ internodal length (0.691), 3^rd^ internodal length (0.767), 4^th^ internodal length (0.452) and thousand grain weight (0.647) (Table 5a). Panicle length had registered high and significant positive correlation with 2^nd^ internodal length (0.477), 3^rd^ internodal length (0.483), 4^th^ internodal length (0.568) and single plant yield (0.505). Primary culm length had a positive relationship with secondary culm length (0.995), 1^st^ internodal length (0.697), 2^nd^ internodal length (0.692), 3^rd^ internodal length (0.790), 4^th^ internodal length (0.383) and thousand grain weight (0.627). Secondary culm length had registered significant positive correlation with 1^st^ internodal length (0.683), 2^nd^ internodal length (0.730), 3^rd^ internodal length (0.793), 4^th^ internodal length (0.366) and thousand grain weight (0.617). The trait 2^nd^ internodal length had recorded the significant positive correlation with 3^rd^ internodal length (0.816), 4^th^ internodal length (0.517) and thousand grain weight (0.588) whereas it has negative association with a number of grains per panicle. The character 3^rd^ internodal length had high and significantly positively correlated with 4^th^ internodal length (0.719) and thousand grain weight (0.508). The trait 4^th^ internodal length was positively with thousand grain weight (0.399) and single plant yield (0.492). Among the quality traits studied, KLBC had high and significant positive correlation with KBBC (0.614), L/B BC (0.757), KLAC (0.633) and L/B AC (0.407). KBBC had registered significant positive and negative correlation with KLAC (0.365) and BER (−0.600), respectively (Table 5b). L/B BC ratio was positively correlated with KLAC (0.501), amylose content (0.426) and negatively correlated with single plant yield (−0.441). KLAC had a high and positive association with the traits namely L/B AC (0.820) and LER (0.650), respectively. KBAC had a significant positive correlation with BER (0.617) and GC (0.512). L/B AC had registered positive and negative significant association with LER (0.412) and BER (−0.550). The character breadth wise expansion ratio exhibited significantly positively correlated with gel consistency (0.442).

**Table 5a.** Genotypic correlation coefficients among different quantitative traits of selected mutants (Semi-dwarf and early mutants) in M3 generation (gamma rays) of improved white ponni

**Table 5b.** Genotypic correlation coefficient among different quality traits of selected mutants (Semi-dwarf and early mutants) in M3 generation (gamma rays) of improved white ponni

### Molecular characterization

#### Polymorphism information content (PIC)

Out of 154 mutants, 30 unique mutants (11 confirmed semi-dwarf and early mutants, 19 early mutants) were selected and subjected to molecular analysis in an M_3_ generation. Fifty-six SSR primer pairs were used, which includes three gene-specific markers for the *sd1* locus. Out of these 56 SSR primer pairs, 45 primers exhibited polymorphism between the mutants and found that a total of 96 alleles (Table 6a) (Fig 4). Alleles per locus ranged from 1 to 3 with a mean of 2.13. It was found that PIC value ranges from 0.037 (RM246) to 0.949 (GA20 Oxi_2) with an average of 0.382. There were several markers which recorded high PIC value and they are highly polymorphic in nature. The markers *viz*., GA20 Oxi_2 (0.949), RM310 (0.662), RM_1_40 (0.648), RM7365 (0.633), RM3431 (0.616) had recorded PIC value more than 0.6. Several other markers namely GA20 Oxi_3 (0.584), RM25 (0.570), RM3555 (0.537), RM493 (0.531), RM5720 (0.529), RM240 (0.523) and RM3912 (0.513) had registered PIC value more than 0.5.

**Table 6a.** Allelic distribution and PIC values of SSR markers in this study

### Marker-trait associations

Single marker analysis was also done and out of 45 polymorphic SSR primer pairs, only 25 primers had a significant correlation with traits studied (Table 6b). An amount of phenotypic variation (R^2^) accounted by markers were estimated and this clearly explains the existence of phenotypic variation to the total variance. RM 310 showed the maximum phenotypic variance of 65.20 per cent and associated with the trait primary culm length whereas the marker RM7365 explained minimum phenotypic variance and linked with a gel consistency. The marker associated with the trait days to 50% flowering was RM335 and RM3452 with an R^2^ value of 37.30 and 31.30 per cent. The marker RM 252 was highly associated with the traits plant height and 1^st^ internodal length with the R^2^ value of 62.40 and 49.20 per cent and several other markers were also associated. The marker RM310 was highly associated with the traits primary culm length, secondary culm length, 4^th^ internodal length, thousand grain weight, KLBC, L/B BC and LER. The marker RM 3912 recorded high association with the traits 2^nd^ and 3rd internodal length with high R^2^ value. The marker RM587 was highly associated with the traits number of grains and KLAC with the R^2^ value of 36.20 and 37.30 per cent. The marker RM167 exhibited a high association with the trait number of productive tillers with the R^2^ value of 21.80 per cent. The marker RM7365 had exhibited a highly significant correlation with the kernel breadth before cooking and explains 21.10 per cent variation to the total variance. The marker RM3431 showed significant association with the traits KBAC and BER with the variability percentage of 22.70 and 30.30 per cent to the total variance. The markers RM551, RM38 and RM246 recorded significantly associated with the traits L/B ratio after cooking, amylose content and gel consistency.

**Table 6b.** Marker-Trait Associations estimated by simple linear regression analysis (SLRA)

### The GA_3_ response of dwarf and early mutants using the micro-drop method

Second leaf sheath length was measured in all identified dwarf mutants after the application of GA_3_ to the coleoptiles of the seeds after germination (Table 7). The GA_3_ response was estimated based on the length of second leaf sheath of GA_3_ treated and non-treated seedlings as a control. The GA_3_ response was estimated on two different durations of 5^th^ day and 15^th^ after treatment. The mean 2^nd^ leaf sheath length of mutants on 5^th^ DAT in non-treated seedlings was 1.60 with the range of 0.92 (IWP E-4) to 1.93 (IWP 48-2) whereas in GA_3_ treated seedlings it was ranged from 1.52 (IWP E-4) to 2.98 (IWP 59-1). On 15^th^ DAT, the mean second leaf sheath length of mutants in non-treated seedling was ranged from 2.30 (IWP 1-12) to 3.03 (IWP 48-2) whereas in GA_3_ treated seedlings it was ranged from 2.98 (IWP 7-1) to 3.89 (IWP 43-1, IWP 33-2). The maximum GA_3_ response was exhibited by the mutant IWP 18-1 (189.80) and minimum GA_3_ response had recorded by the mutant IWP 50-11 (112.26) on 5^th^ DAT whereas in 15^th^ DAT the mutant IWP 18-1 (146.84) recorded the maximum response to GA_3_ application and mutant IWP 48-2 (104.29) had registered the minimum response. The mutants IWP 47-2 (120.23), IWP 11-2 (134.04), IWP 1-1 (139.48), IWP 48-2 (122.45), IWP 7-1 (133.39), IWP 1-12 (140.35) and IWP 50-11 (112.26) had recorded less GA_3_ response when compared wildtype (149.12) on 5^th^ DAT. On 15^th^ DAT, the mutants *viz*, IWP 47-2 (107.44), IWP 59-1 (117.82), IWP 1-1 (120.38), IWP 48-2 (104.29), IWP 7-1 (110.78) and IWP 50-11 (117.10) exhibited less GA_3_ response when compared to wildtype genotype (159.69). In 5^th^ DAT, the maximum and minimum shoot length of non-treated seedlings was recorded by the mutant IWP 43-1 (10.40 cm) and IWP E-4 (3.60 cm) whereas in treated seedlings the maximum and minimum shoot length was exhibited by the mutants IWP 1-12 (14.50 cm) and NLM (7.67 cm) (Table 8) (Fig 5). In 15^th^ DAT, shoot length of non-treated seedlings of mutants were ranged from 4.60 cm (IWP 18-1) to 12.00 cm (IWP 43-1) whereas the shoot length of treated seedlings was ranged from 10.73 cm (IWP 50-11) to 21.33 cm (IWP 48-3).

**Table 7.** GA3 response of dwarf and early mutants using the microdrop method

**Table 8.** Shoot length variation after application of GA3 by the microdrop method Figures

### α – amylase activity

The identified dwarf and early mutants of Improved White Ponni were subjected to study α – amylase activity induced by the application of GA_3_ induction by GA application. In this study, α – amylase production and secretion were observed as plaques in the wild-type as well as in dwarf and early mutants *viz*., IWP 43-1, IWP 47-2, IWP 59-1, IWP D-1, IWP 18-1, IWP E-3 and IWP E-4. Based on the staining pattern of the starch plate with iodine solution revealed that the mutants namely IWP 50-11 and IWP 33-2 showed there is no α – amylase production which indicated that these two mutants were related to the GA pathway (Fig 6). The other mutants *viz*., IWP 43-1, IWP 47-2 and IWP 18-1 exhibited α – amylase production was significantly decreased when compared to wild type. The mutants IWP 59-1, IWP D-1, IWP E-3 and IWP E-4 were produced more α – amylase than wild-type.

### Study on cell arrangement pattern in unique mutants of Improved White Ponni

Internodal regions of dwarf and early mutants and leaf sections of identified (IWP 59-1) and of Improved White Ponni along with wild-type were subjected to observe under scanning electron microscope to study the pattern of internal cell arrangements. Cell patterning differences were observed and revealed that the semi-dwarf mutant of White Ponni exhibited larger cell size and less number of cells arrangement per unit area than the wild type (Fig 7).

## Discussion

Micro-mutational events with the least deleterious effects were considered to be the most reliable and effective mutations in providing variability for quantitative traits. The mutagens which offered to induce variability is utmost important for any crop breeding program since events created by mutation may alter some large or small phenotypic expression. Besides the observation of frequency of chlorophyll mutations across the seedlings of M_2_ generations, changes in quantitative traits were also observed in both M_2_ and M_3_ generations. Gaul (1964) classified viable mutations as macro and micro mutations, while Swaminathan (1965) grouped them as macromutations and systematic mutations. Besides the usage of chlorophyll mutations observed in Improved White Ponni to estimate the effectiveness and efficiency of mutagen, viable mutants for plant type *viz*., early/late flowering, dwarf, high tillering habit, narrow rolled leaf, upper albino, lodging resistant, grassy and extreme dwarf and lanky culms were observed in M_2_ population. In case of plant type modifications, semi-dwarf and narrowly rolled leaf mutants appeared in mutated population. The frequency of semi-dwarf mutants was high in lower doses of gamma radiation when compared to higher doses in gamma-irradiated populations. Shadakshari *et al.* (2001) reported the occurrence of higher frequency of dwarf/semi-dwarf non-lodging early flowering and high yielding mutants in five rice varieties treated with gamma rays. Yankulav *et al.* (1980) and Reddi and Rao (1988), who reported that reduction on mutagen’s effectiveness were higher with increase in their dosage and our study also reported the same. Apart from estimating the mutagen efficacy and efficiency, chlorophyll and viable mutation frequency estimation become useful for selecting the suitable mutagen in crop breeding (Nilan et al., 1965).

The viable mutations isolated in the study showed changes in major traits which could be utilized in the future breeding programme where reshuffling of traits may be tried by conventional breeding methods. However, most of the morphological mutants identified in M_2_ generation failed to inherit in M_3_ generation. According to the statements of Luo *et al.* (2012), these characters may be controlled by recessive genes or are susceptible to the environment. Moreover, whatever the changes occurred in the plants due to mutation is an error according to the plant’s geometry. They tend to rectify it in due course through recombinational events. That is why most of the observed mutants were not inherited in future generations. Thus, evolving a new phenotype with consistent expression through mutation is a chance event rather than a choice.

### Induced variability on quantitative traits in M_2_ and M_3_ generations

In the present investigation, comparison on means and variances (phenotypic and genotypic) of various quantitative traits including the quality parameters in M_2_ and M_3_ generations indicated a considerable shift in mean and variability in the treatments and this could be because of recombination happened in the M_1_ plants. Siddiqui and Singh (2010), who found out that the variability may increase significantly with respect to increase or decrease in the mean values of the trait studied. Our results were also accorded with earlier reports and this may be expected due to the reason that, recombinational events due to mutation creates more variations in the living systems (Johnston, 2001).

The rate at which different mutations occurring at a specified stage is mutation rate and this estimate can be made when the targeted mutation is very obvious and detectable. But most of the quantitative traits do not follow this pattern and making the situation of detecting mutations very complex. Under these circumstances, mutational heritability [genetic variance increase by a single generation of mutation (*Vm*)/the environmental variance of the trait (*Ve*)) is estimated (Houle *et al*, 1996) and one has to measure the amount of new genetic variation arising in each generation due to mutation. Moreover, the variations for quantitative traits observed in the M_1_ generation have no significant importance in deciding the number of mutations for the targeted trait. The detection of mutations for a quantitative trait has to be decided based on the progeny values *i.e*. M_2_ generation. The mutant population which exhibits high mean coupled with a high variance for a trait is the first choice of selection. Usually, mutations are detected when there is huge phenotypic effect on a trait has been observed in large mutant population of a variety or elite cultivar.

Investigation on M_2_ generation had shown higher variation in 100 Gy for the trait days to fifty per cent flowering. The genetic parameters were very low in M_3_ generation when compared to M_2_ generation treated with gamma rays. Nayudu *et al.* (2007) and Anilkumar (2008) had also reported similar findings of our study. The trait plant height showed wider differences of all genetic parameters and the selection was effective in M_2_ and M_3_ as reported by Singh *et al.* (2006). The traits namely panicle length, productive tillers and grains per panicle shown moderate to high variability in M_2_ and M_3_ indicating the scope for selection and improvement (Kumar *et al.*, 2006). These traits expresses moderate to high h^2^ and low to high GA in M_2_ and M_3_ generation and these results are lined with Ahmed *et al.* (2010). Primary culm length and secondary culm lengths had exhibited moderate variability, moderate to high h^2^ and low to moderate GA (Bin-mei *et al.*, 2006). A similar trend was also observed in M_3_ generation for the traits *viz*., 1^st^ internodal length, 2^nd^ internodal length, 3^rd^ internodal length and 4^th^ internodal length. Variability study clearly revealed that these traits showed a slightly wider variation in the mutant population (Zou *et al.*, 2005; Bin-mei *et al.*, 2006). Thousand grain weight and single plant yield exhibited minimum variability, low to high h^2^ and GA in M_2_ and M_3_ generations (Kishor *et al.*, 2008; Yadav *et al.*, 2010). These two parameters (h^2^ and GA) was a trustworthy real estimate ifor selection than heritability alone. Conversely, it is not essential criteria that a trait conveying high heritability will also display high genetic advance. This is because; the heritability calculation leads to estimation errors and largely depends on genotype-environment interactions. Estimation of the coefficient of variation specifies only the available variability of the particular trait and it does not provide any information about heritability.

Genetic parameters for different quantitative traits were estimated and most of the traits exhibited varied and significant mean performance with a high level of variation among the mutants. The traits namely, plant height, days to 50 % flowering, panicle length, productive tillers per plant, grains per panicle, thousand grain weight and grain yield per plant exhibited slightly wider variability, higher h^2^ and moderate to high GA among the mutants. These genetic parameters could be valuable measure for the effective selection towards yield enhancement is possible (Kishor *et al.*, 2008; Yadav *et al.*, 2010; Akinwale *et al.*, 2011). Among the grain quality traits studied, most of the traits exhibited higher variation among the mutants and some of the putative mutants had on par quality with the wildtype. The results were accorded with the findings of Siddiqui and Sanjeeva (2010) and Subbaiah (2011). In this study, major emphasis and attention were given to the selection of dwarf and early mutants, which determines crops per year and adaptation towards maximizing the yield over seasonal and regional specification. The identified semi-dwarf and early mutants had shown approximately 35 to 45 per cent reduction in plant height and 4 to 33 days earlier in duration than wildtype.

The magnitude of association of component traits assists the plant breeder to improve the yield and other important traits. Among the biometric traits studied, the traits panicle length and 4^th^ internodal length had a significant positive relationship with grain yield as reported as well (Sankar *et al.*, 2006; Anilkumar, 2008; Immanuel *et al.*, 2011; Bagheri *et al.*, 2011;Akinwale *et al.*, 2011). Saif-ur-Rasheed *et al.* (2002a) reported the association of a number of tillers and productive tillers had an optimistic relationship with grain yield. The trait plant height was positively associated with the traits panicle length, primary culm length, secondary culm length, 1^st^ internodal length, 2^nd^ internodal length, 3^rd^ internodal length, 4^th^ internodal length and thousand grain weight. The selection based on the above traits highly influences the grain yield of the mutants and provides an indirect selection of these traits would ultimately increase the yield quantum (Rashid *et al.*, 2013). Another trait primary culm length exhibited positive and significant correlation with the traits secondary culm length, 1^st^ internodal length, 2^nd^ internodal length, 3^rd^ internodal length, 4^th^ internodal length and thousand grain weight. Another trait secondary culm length exhibited positive and significant correlation with the traits 1^st^ internodal length, 2^nd^ internodal length, 3^rd^ internodal length, 4^th^ internodal length and thousand grain weight as reported by Muhammad *et al.* (1982). The trait 4^th^ internodal length had registered significant positive association with thousand grain weight. Apart from morphological traits, quality parameters of the mutants were also shown on par in most of the traits observed with the positive relationship among them and these mutants would be resulted to improve those traits through effective breeding strategies (Khatun *et al.*, 2003; Veni *et al.*, 2003).

### Molecular characterization and Marker-trait Associations

Semi-dwarf and early mutants, narrowly rolled leaf mutants of Improved White Ponni generated through gamma radiation were identified in M_2_ were forwarded to M_3_ (30 mutants) generations. These mutants were surveyed using 55 microsatellite markers to detect the polymorphism among mutants and it was noticed that 96 alleles were detected. Among the SSR markers, RM_2_46 and GA20 Oxi_2 found to be have lower and higher PIC value of 0.037 and 0.949 with a mean value of 0.382. The microsatellites *viz*., GA20 Oxi_2, RM310, RM_1_40, RM7365, RM3431, GA20 Oxi_3, RM_2_5, RM3555, RM493, RM5720, RM_2_40 and RM3912 had recorded PIC value more than 0.5 (Chen *et al.*, 2011; Kumar and Bhagwat, 2013). The PIC value made the reflection of allelic diversity between the mutants and not found to be uniformly high for the SSR loci tested (Wang *et al.*, 2009; Pervaiz *et al.*, 2010). These findings were supportive that, microsatellites are more potent and informative to study the genetic divergence and variability pattern of closely related individuals (Xu *et al.*, 2004).

Marker data were also correlated with traits studied in rice mutants for determining the informative SSR markers associated with these traits. Out of 45 polymorphic SSR primer pairs, only 25 primers had a significant association with different traits studied. Single marker analysis exhibited that, the markers RM3912 and RM430 was highly associated with the trait plant height. The marker RM302 had exhibited a high association with the traits plant height, primary culm length, secondary culm length, 1^st^ internodal length and 4^th^ internodal length. The marker RM302 on chromosome 1 had shown putative linkage with semi-dwarfism (Subashri *et al.*, 2008, Wang *et al.*, 2009). Another gene (*sd-1*) based marker, GA20Oxi_2 was positioned near to the marker was also found linked with semi-dwarf trait. Asano *et al.* (2007) also reported that the dwarfism is due to complete loss of *sd-1* gene function. The marker RM310 showed association with primary culm length, secondary culm length, 1^st^ internodal length, 2^nd^ internodal length, and 3^rd^ internodal length. Several grain-quality traits were observed in mutants selected from M_3_ generation of White Ponni and marker-trait association trait study revealed that the marker RM7365 had exhibited a highly significant correlation with the KBBC, the marker RM3431 showed significant association with the traits KBAC and BER. The markers RM551, RM38 and RM246 recorded significantly associated with the traits L/B ratio AC, amylose and GC. Roy *et al.* (2006) reported that an association study in wheat showed a total of 99 of the 221 polymorphic SSR bands and explained a maximum of R^2^ value of 8.12 per cent for tiller numbers to 27.95 per cent for harvest index (Monfared *et al.*, 2008; Kalivas *et al.*, 2011).

### The response of mutants to GA_3_ application

GA_3_ responses of semi-dwarf and early mutants were determined using a micro-drop method by GA_3_ application on the coleoptiles region of the seeds after three days of germination. In the present investigation of GA_3_ response, all the mutants were recorded moderate to high response to the GA_3_ application. Narrowly rolled leaf mutant also recorded low response to the GA_3_ application (Ueguchi-Tanaka *et al.*, 2000). The shoot length was also observed in all the mutants five and fifteen days after GA_3_ application which revealed that the increase in shoot length of semi-dwarf and early mutants showed variation in their responsiveness. Thus, this study clearly emphasized that all the mutants identified as GA_3_ response mutants except IWP 11-2, IWP 48-2, IWP 50-11 and IWP 33-2 which showed a very minimal response to the GA_3_ application after fifteen days and these were GA_3_ insensitive mutants. Ogi *et al.* (1993) discovered that the *sd-1* gene reduced the internode length by preventing cell division in internodes. Other researchers have proved that the *sd-1* gene was sensitive to GA_3_, which influenced the content of GA_3_ and the vigour of peroxidase in the plant (Shi and Shen, 1996; Gao *et al.*, 2009 and Asano *et al.*, 2009). Application of GA_3_ in ig*a-1* semi-dwarf mutant resulted in the partial restoration of wildtype plant height and this was confirmed as insensitive and similar result was also reported by the current study, were both sensitive and insensitive mutants were identified (Wang et al., 2009).

### α – amylase activity of semi-dwarf and early mutants

The α-amylase production in assay plates and shoot elongation by the exogenous application of GA_3_ are GA-mediated process (Matsukura *et al.*, 1998). The staining pattern of agar plate revealed that the mutant’s *viz*., IWP 50-11, IWP 33-2, IWP 43-1, IWP 47-2 and IWP 18-1 had exhibited low production of α- amylase which ultimately infers that these mutants have some defects in the GA related pathway (Qin *et al.*, 2008). The secretion of α-amylase was found to be noticed as low around the seed placed in the assay plates containing GA_3_ in both wildtype and mutants and this was lined with the earlier report in the d62 mutant (Li *et al.*, 2010). On overall conclusions, it was found that the identified semi-dwarf early mutants were either GA_3_ responsive or non-responsive in nature (Lanahan and Ho, 1988; Chandler, 1988 Zou *et al.*, 2005).

### Cell structural pattern analysis in internodal regions of Semi-dwarf mutants

Intercalary meristem cell division and elongation are the major cause for internodal elongation in rice and flaw in these processes severely affects the plant height. A longitudinal section from the internodal regions of the semi-dwarf mutants and wild-type was examined under a scanning electron microscope (SEM) to analyse the inter-cell arrangement patterns. This study revealed that the semi-dwarf mutant exhibited larger cell size (more length and breadth) and a lesser number of cells arrangement per unit area than the wildtype. These results ultimately highlighted that these mutants are defective in GA mediated pathway and therefore cell division was minimized which would ultimately result in the reduced internodal lengths (Wang *et al.*, 2009).

## Conclusion

The overall study concluded that the semi-dwarf mutants identified had a significant reduction in the 2^nd^, 3^rd^ and 4^th^ internodal regions and thereby reduces the plant height (up to 50 per cent reduction), imparted lodging resistance and provided significant yield and quality improvement over the wild-type. The identified early mutants of White Ponni exhibited 4 to 33 days of earliness than the wildtype and thereby aiding to reduce crop duration. The molecular study on these mutants using markers linked to plant height like RM302, GA200xi_1 and GA200xi_2 on chromosome 1 had allelic variations between the mutants. GA200xi_2 showed null alleles in the dwarf mutants and this clearly emphasized that there are some base deletions occurred in the region of exon 2 of *sd-1* region and it can be further studied for the expression profiling. GA_3_ response and α-amylase activity were studied which reported that the identified semi-dwarf mutants were more or less sensitive to GA_3_. These desired mutants of White Ponni might have the potential for further improvement of rice production through future breeding programmes.

## Acknowledgement

The authors thank the BARC (***Board of Research in Nuclear Sciences***) ***BARC, Mumbai*** for financial support.

## Author Contributions

MAP has designed the research plan. RC, APL and MAP conducted the research. RC, APL, KKV and YJK analyzed and interpreted the data. RC wrote the manuscript. The authors approved the final version of the manuscript.

---

Figure 3: Variation in internodal lengths of Improved White Ponni mutants compared to wildtype

Figure 4: Genotypic data of novel Improved White Ponni mutants using microsatellite markers Genotyping result of RM252 Genotyping result of RM310

Figure 5: Variation on Shoot length in early semi-dwarf mutants of Improved White Ponni with and without GA3 application by micro-drop method

Figure 6: Variation in the alpha-amylase activity of unique semi-dwarf mutant in this study a) Wildtype-Improved White Ponni b) Identified mutants. -GA_3_ – without GA_3_ application and +GA_3_ - with GA_3_ application

Figure 7: Variation in cell structural pattern of the third internodal region in Improved White Ponni and Semi-dwarf Mutant (IWP 59-1). This image explaining the cell size (length and breadth of each cell between wildtype and mutant (green markings)

